# *Llamanade*: an open-source computational pipeline for robust nanobody humanization

**DOI:** 10.1101/2021.08.03.455003

**Authors:** Zhe Sang, Yufei Xiang, Ivet Bahar, Yi Shi

**Affiliations:** Department of Cell Biology, Pittsburgh, PA, USA; Department Computational and Systems Biology, Pittsburgh, PA, USA; University of Pittsburgh–Carnegie Mellon University Program in Computational Biology, Pittsburgh, PA, USA

**Keywords:** VHH single-domain antibodies, nanobodies, antibody therapy, SARS-CoV-2, humanization

## Abstract

Nanobodies (Nbs) have recently emerged as a promising class of antibody fragments for biomedical and therapeutic applications. Despite having marked physicochemical properties, Nbs are derived from camelids and may require “humanization” to improve translational potentials for clinical trials. Here we have systematically analyzed the sequence and structural properties of Nbs based on NGS (next-generation sequencing) databases and high-resolution structures. Our analysis reveals substantial framework diversities and underscores the key differences between Nbs and human Immunoglobulin G (IgG) antibodies. We identified conserved residues that may contribute to enhanced solubility, structural stability, and antigen-binding, providing insights into Nb humanization. Based on big data analysis, we developed “*Llamanade*’’, a user-friendly, open-source to facilitate rational humanization of Nbs. Using Nb sequence as input, *Llamanade* provides information on the sequence features, model structures, and optimizes solutions to humanize Nbs. The full analysis for a given Nb takes less than a minute on a local computer. To demonstrate the robustness of this tool, we applied it to successfully humanize a cohort of structurally diverse and highly potent SARS-CoV-2 neutralizing Nbs. *Llamanade* is freely available and will be easily accessible on a web server to support the development of a rapidly expanding repertoire of therapeutic Nbs into safe and effective trials.

**Author Summary:** Camelid Nbs are characterized by small size, excellent pharmacological properties and high flexibility in bioengineering for therapeutic development. However, Nbs are “xeno” antibodies, which require “humanization” to improve their translational potential. Currently, there is a lack of systematic investigation of Nbs to rationally guide humanization. No dedicated software has been developed for this purpose. Here, we report the development of *Llamanade*, an open-source computational pipeline and the first dedicated software to facilitate rational humanization of Nbs.

To subjectively evaluate *Llamanade*, we used it to humanize a cohort of structurally diverse and ultrapotent antiviral Nbs against SARS-CoV-2. Robust humanization by *Llamanade* significantly improved the humanness level of Nbs to closely resemble fully human IgGs. Importantly, these highly humanized antiviral Nbs remained excellent solubility and comparably high bioactivities to the non-humanized Nb precursors. We envision that *Llamanade* will help advance Nb research into therapeutic development.

## Introduction

V_H_H antibodies or nanobodies (Nbs) are small antigen-binding fragments that are derived from camelid (e.g., llama, alpaca, dromedary, and camel) heavy-chain antibodies (1). Nbs are composed of four conserved framework regions (FRs) that fold into β-sandwich core structures (2). Three hypervariable loops, or complementarity-determining regions (CDRs), are supported by robust scaffolds to provide antigen-binding specificity. It has been shown that Nbs may preferentially target concave epitopes to efficiently interact with target antigens. In many cases, the binding is markedly different from heterodimeric immunoglobulin G (IgG) antibodies, where the epitopes are generally more flat or convex (3-5).

The small size (∼ 15 kDa), robust fold, and lack of glycosylation enable rapid production of Nbs in microbes at low costs. Affinity matured Nbs are characterized by excellent physicochemical properties including high solubility and stability, which are critical for drug development, production, transportation, and storage. While Nbs are monomeric, they can be easily bioengineered into bispecific and multivalent modalities to achieve avidity binding, which may resist the mutational escape of the target (e.g., virus and cancer antigen) under selection pressure, and/or incorporate additional new functionalities (6-8). Because of their small size, Nbs can bind compact molecular structures and may penetrate tissues more efficiently than large IgG antibodies, thus facilitating molecular and diagnostic imaging applications (9, 10).

In response to the COVID-19 (Coronavirus disease 2019) pandemic, thousands of highly potent and neutralizing Nbs have recently been identified by using *in vivo* affinity maturation coupled to a robust Nb drug discovery pipeline (8). These multi-epitope Nbs specifically target the receptor-binding domain (RBD) of SARS-CoV-2 spike glycoprotein with high affinity, and are cost-effective antiviral agents for the evolving virus(5). The outstanding preclinical efficacy of an inhalable construct has been recently demonstrated for inhalation therapy of SARS-CoV-2 infection by small Nb aerosols (11). At an ultra-low dose, this innovative therapy has been shown to reduce lung viral titers by 6-logs to minimize lung pathology and prevent viral pneumonia (11). Moreover, high-resolution structural analyses have facilitated epitope mapping and classification of potent neutralizing Nbs into three main classes, which are characterized by distinct antiviral mechanisms. Systematic structural studies have provided insights into how Nbs uniquely target the spike to achieve ultrahigh-affinity binding and broadly neutralizing activities against SARS-CoV-2 and its circulating variants (5).

Owing to these unique properties, Nbs have emerged as a compelling class of biologics (12). The first Nb drug (Cablivi) has recently been approved by the US Food and Drug Administration (FDA) (13); more candidates are undergoing clinical trials (4). While these efforts have greatly inspired the innovative medical uses of Nbs and antibody fragments, there are remaining challenges for safe and effective applications to diseases in humans. In particular, anti-drug antibody (ADA) responses can reduce drug efficacy and, in rare cases, cause exacerbated inflammatory responses and toxicity (14). The underlying mechanism of ADA remains to be fully understood, however, several factors including, critically, the use of non-human antibodies may contribute to the side effects (15). The consensus is that the humanization of xeno-species antibodies is necessary for drug development. Here “humanization” refers to increasing the similarity of antibodies of non-human origins to human antibodies. Thus far, efforts to humanize non-human antibodies has been exemplified by the clinical benefits of humanizing murine antibodies (16). Humanized and fully human IgGs now dominate clinical development of biologicals (17).

Similar to the humanization of murine antibodies, strategies for Nb humanization are based on CDR grafting or FR resurfacing. One strategy involves 1) grafting antigen-specific CDRs to a specific human heavy chain variable domain (VH_human_) framework, which often is a universal Nb framework (18). While this method has been successfully applied to some Nbs, using a single framework as the scaffold template may undermine the structural compatibility with many CDRs. While generally conserved, antibody frameworks nevertheless show substantial sequence and structural diversity to support infinite CDR loop conformations for antigen recognition. Such high scaffold diversity can not be fully represented by a small number of germline sequences. Another strategy centers around 2) resurfacing solvent-exposed frameworks without changing buried residues with the guidance of available structures or structural models. Resurfacing is based on the assumption that solvent-exposed, non-human residues are less likely to contribute to the structural integrity and/or antigen engagement. In addition, unique CDR properties of Nbs, which remain to be fully investigated, may also contribute to the development of ADA. Overall, there is a lack of systematic and structural investigations into Nb humanization, which is critical for moving therapeutic Nbs into clinical trials.

In this study, we have leveraged antibody/Nb next-generation sequencing (NGS) datasets and high-resolution structural information from the Protein Data Bank (PDB) to systematically analyze and compare Nbs, and mammalian (specifically, human and murine) IgGs. Our analysis reveals unique sequence and structural properties of Nbs and provides insights into Nb humanization. Guided by the large-scale analysis, we have developed *Llamanade-* an open-source software to facilitate Nb humanization. *Llamanade* can rapidly optimize the solution and provide quantitative measurement of the extent of humanization. Finally, we applied *Llamanade* to humanize a set of structurally diverse and ultrapotent SARS-CoV-2 Nbs. These humanized antiviral Nbs closely resemble VH_human_ frameworks and have demonstrated excellent solubility and bioactivities comparable to the non-humanized precursor Nbs.

## Results

### Systematic and quantitative analysis of Nbs and IgG antibodies

We compiled a high-quality Nb sequence library by randomly selecting 49,686 distinct, natural Nb sequences derived from previously generated NGS databases (19). Non-redundant human and murine IgG sequences (22,450 and 10,696, respectively) from the EMBL-Ig database were also selected to generate IgG sequence libraries (20). The variable region of antibody sequences and Nbs were numbered, and their CDRs and FRs were annotated using the Martin numbering scheme (21). To quantify the “humanness” level of Nbs, we used the T20 score which has been shown to distinguish VH_human_ and VH_mouse_ with high specificity (22). In addition, the T20 score may positively correlate with decreased immunogenicity of therapeutic antibodies to potentially inform clinical studies (22). An input Nb will be searched against our curated human IgG library to obtain the top 20 matched VH_human_s based on sequence identity. The percentage identity of the top 20 matches will be averaged to obtain the T20 score (**Method**). We calculated the T20 scores for the framework region (T20_FR_) of all the Nbs, VH_mouse,_ and VH_human_ in the respective libraries to generate the distribution plots (**Figure 1A, Figure S1**). As expected, the scores of different antibody species largely follow normal distributions with VH_human_ slightly skewed towards the lower end. Compared to VH_mouse_, Nbs are significantly more similar to VH_human_, which is consistent with the previous analysis using a limited number of sequences(23). Notably, a substantial fraction of the Nbs is indistinguishable from VH_human_ based on overlapping T20 scores. The average T20_FR_ for Nbs and VH_mouse,_ are 79.7 and 72.1, respectively (**Figure 1A**). High similarity between Nbs and VH_human_ sequences is primarily attributed to FR1, FR3, and FR4 among the four FRs (**Figure 1B, 1D, 1E**). In contrast, the FR2s of Nbs are markedly different from their human counterparts (**Figure 1A**). Moreover, while framework sequences are generally conserved, substantial variations are also evident (σ_human VH_ = 4.55, σ_mouse VH_ = 4.22, σ_Nb_= 3.73). These structural variations underlie the potential challenges of using a single scaffold template to humanize a large repertoire of highly structurally diverse Nbs.

**Figure 1.**
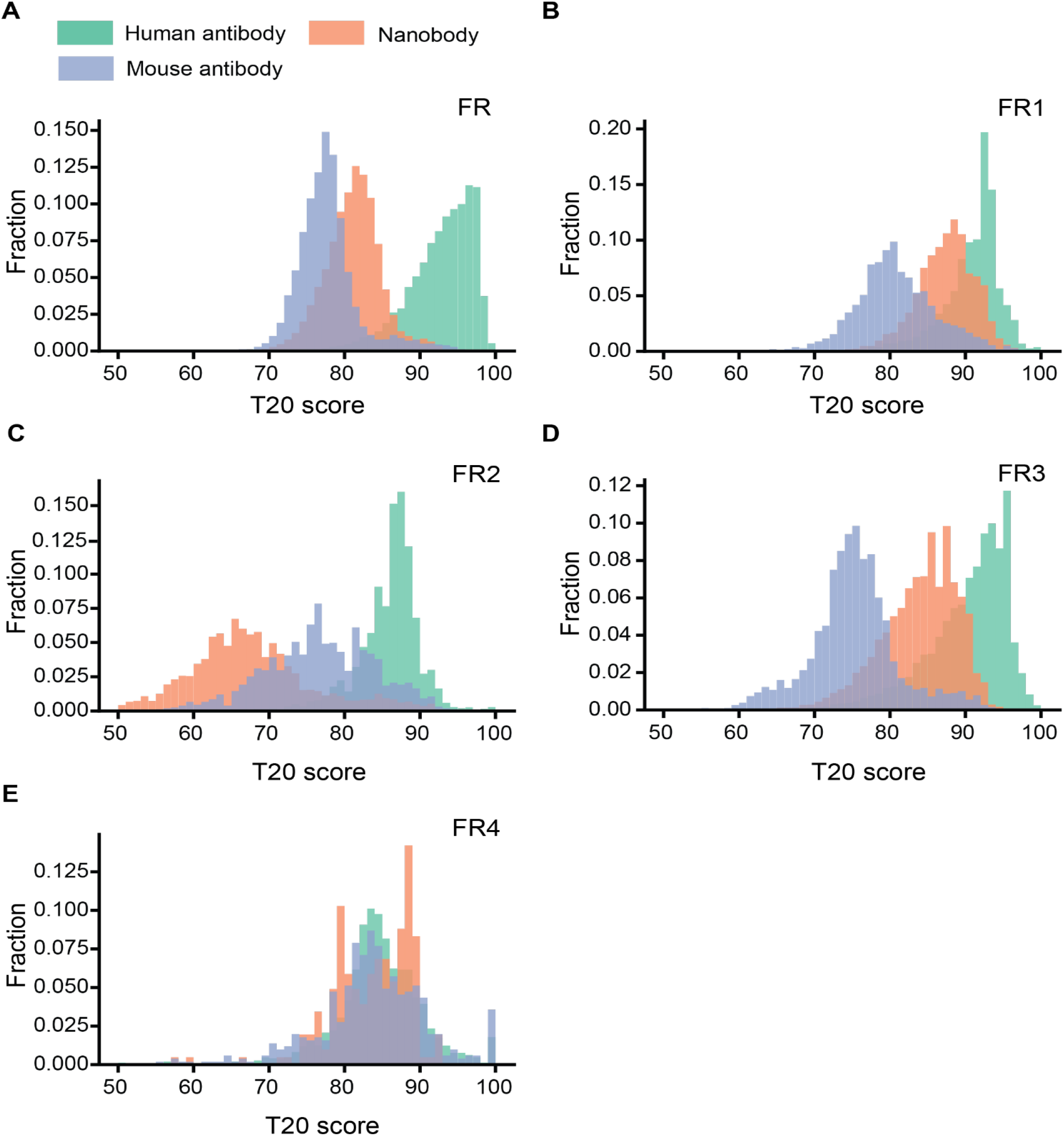
Analysis of T20 humanness scores for the framework sequences of Nbs and VHs. **A-F**. Distributions of the T20 humanness scores based on different FR regions. FR: framework.

Identifying framework sequences that differ between Nbs and the top matched VH_human_ is likely critical to inform humanization strategies. Nbs and the corresponding VH_human_ were aligned and the resulting sequence logo profiles are presented in **Figure 2A-D** (**Methods**). The average percentages of residue substitution are 10%, 12.8%, 28.6% and 4% for the respective four FRs (**Figure 2E**). Hotspot residue substitutions (from Nbs to VH_human_) were identified including FR1 residues: A14P, R27F; FR2 residues: F/Y37V, E44G, R45L, F/L47W; FR3 residues: V75L, K83R, P84A, A94R/K; and FR4 residue: Q108L. While the substitutions on three FRs share similar physicochemical properties, FR2 substitutions, especially F/Y37, E44, and R45, may reduce local hydrophilicity of Nbs. Large-scale sequence analysis reveals key differences between Nbs and VH_human_ which may be modified to achieve higher similarity to human antibodies.

**Figure 2.**
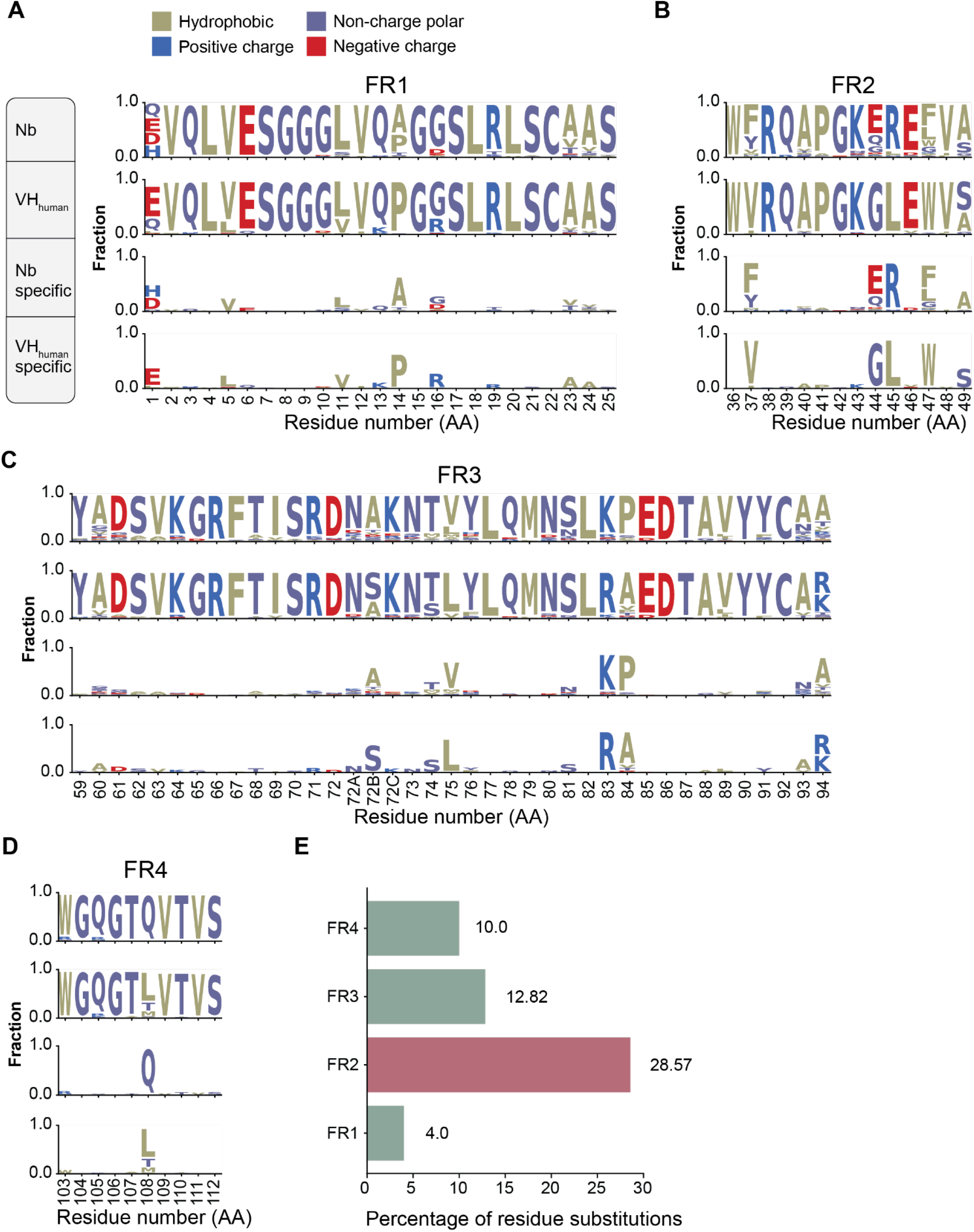
Sequence conservation analysis of Nbs and VH_human_. **A**)-**D**) Sequence logos of framework regions (FR1, FR2, FR3, and FR4) of Nbs and VH_human_. In each panel, the top diagram shows the profile of Nbs, the second shows that of VH_human_, the third shows the conserved residues specific to Nbs and the last, those specific to VH_human_. **E**) Percentage of conserved amino acid substitution on Nb FRs compared to VH_human_.

### Large-scale structural analysis reveals residue interactions specific to Nbs

Given the structural differences of Nbs from VH_human_, indiscriminate substitution of camelid Nb residues for human VH_human_ residues may result in perturbations to the overall Nb structure. In addition, surface exposed residues are more likely to be recognized by the host immune response and are thus of particular interest for humanization. To minimize immunogenicity without perturbing the overall structure, we systematically analyzed high-resolution structures of antigen-antibody interactions to gain insights into the crucial, unique structural properties of Nbs. We assessed the structures of 190 distinct Nbs and 621 human IgGs from the AbBank (20). This structural information was used to differentiate buried residues from surface-exposed residues, which may elicit immunogenicity and therefore require humanization. Here, a residue is considered “buried” if the projecting side chain is 3 Å or more below the antibody surface (24) (**Methods**). As expected, a small fraction of antibody FR residues are buried while the majority are solvent-exposed. Moreover, compared to VH_human_ (average 73%, σ=6.2%), significantly more Nb FR residues (average 76.9%, σ = 6.3%) are solvent-exposed (**Figure 3A, 3B**). The major differences occur in the FR2, where R45 and F/L47 (Nbs) are fully exposed while the corresponding human residues are buried (**Figure 3C, 3D)**. These solvent exposed residues in FR2 are likely to be important for the high solubility of Nbs and are not recommended for humanization.

**Figure 3.**
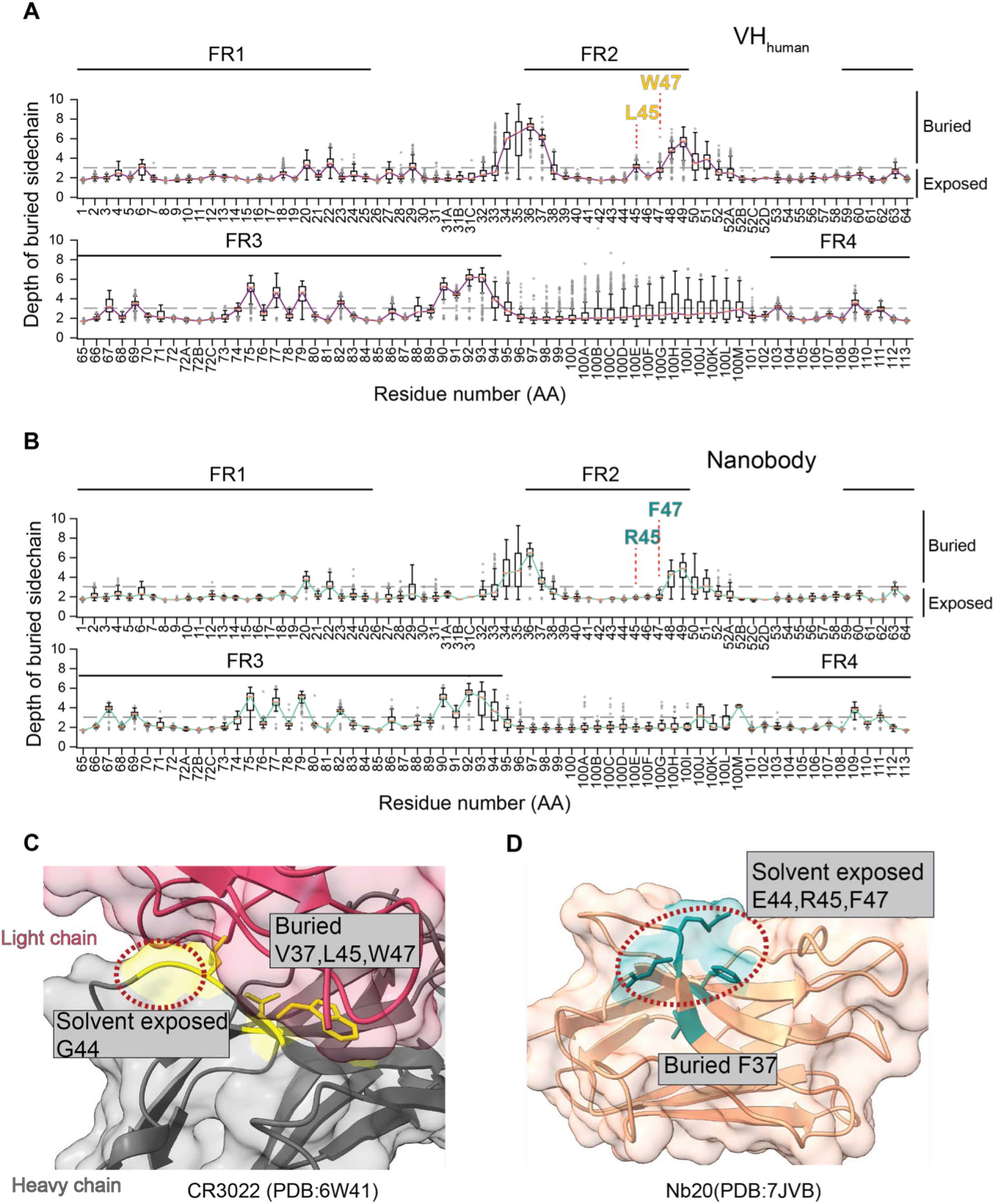
Analysis of buried residues in Nbs and VH_human_. **A**. Distribution of depth of burial of residue side chain at the aligned position of the VH_human_. Semi solvent-exposed FR2 residues L45 and W47 are highlighted. Error bars represent standard deviations. **B**. Same as **A** for the aligned position of Nbs. Fully solvent-exposed FR2 residues R45 and F47 are highlighted. **C**. Ribbon diagram of the human antibody illustrating the extent of burial of FR2 residues (PDB: 6W41). The light chain is colored pink and the heavy chain is gray. Among 4 hallmark FR2 residues (37, 44, 45, and 477), only G44 is solvent-exposed. **D**. Nb ribbon diagram showing burial of FR2 residues (PDB: 7JVB). Among 4 hallmark FR2 residues, E44, R45, and F47 are solvent-exposed.

Large-scale structural analysis also reveals two types of intramolecular interactions that may contribute to structural integrity. The first is between FR2 and FR4. Here, W103 (FR4) can associate with R45 (FR2) by a conserved cation-π interaction, which is replaced by hydrophobic interaction of L45: W103 in VH_human_. In addition, W103 can also interact with F/Y37 using a π-π interaction, which is absent in the IgG VH domains. To substantiate these observations, we measured the distances and angles of the charged atoms of R45 with respect to the centroids of aromatic rings from W103. We also measured those between F/Y37 and W103 (**Method**). The average distance/angle of the R45:W103 and F/Y37:W103 interactions are 3.9 Å/65.7°, and 4.1 Å/31°, supporting the respective bond formations (cation-π and π-π interactions) (25, 26). These interactions should be conserved as they may be critical to support the robust scaffold and excellent physicochemical properties of Nbs.

The second important intramolecular interaction specific to Nbs is between the FR2 and CDR3 (**Figure 4D**). The frequency of this bond positively correlates with CDR3 length. Over 80% of long CDR3s (> 15 residues) were found to have this interaction (**Figure 4D**). In addition, most long CDR3’s contain a small helix structure (typically 3-4 amino acids in length) that was rarely identified on VH_human_ (**Figure 4E, 4G**). Analysis reveals that long CDR3 loops of Nbs can fold back to interact with two specific FR2 residues (positions 37 and 47, **Figure 4F**), where the corresponding VH_human_ residues use to interact with the light chains (VLs) (27) (**Figure S2A**). This FR2: CDR3 interaction may impact Nb humanization in two ways: 1) the long CDR3 may shield potentially immunogenic residues on FR2, which are otherwise solvent-exposed. 2) The helix associated with long CDR3 loops may potentially enhance immunogenicity and the development of ADA in clinical applications (**Figure 4G**). Nevertheless, these FR2 residues unique to Nbs must be conserved, as resurfacing these residues may alter CDR3 conformations and compromise antigen binding.

**Figure 4.**
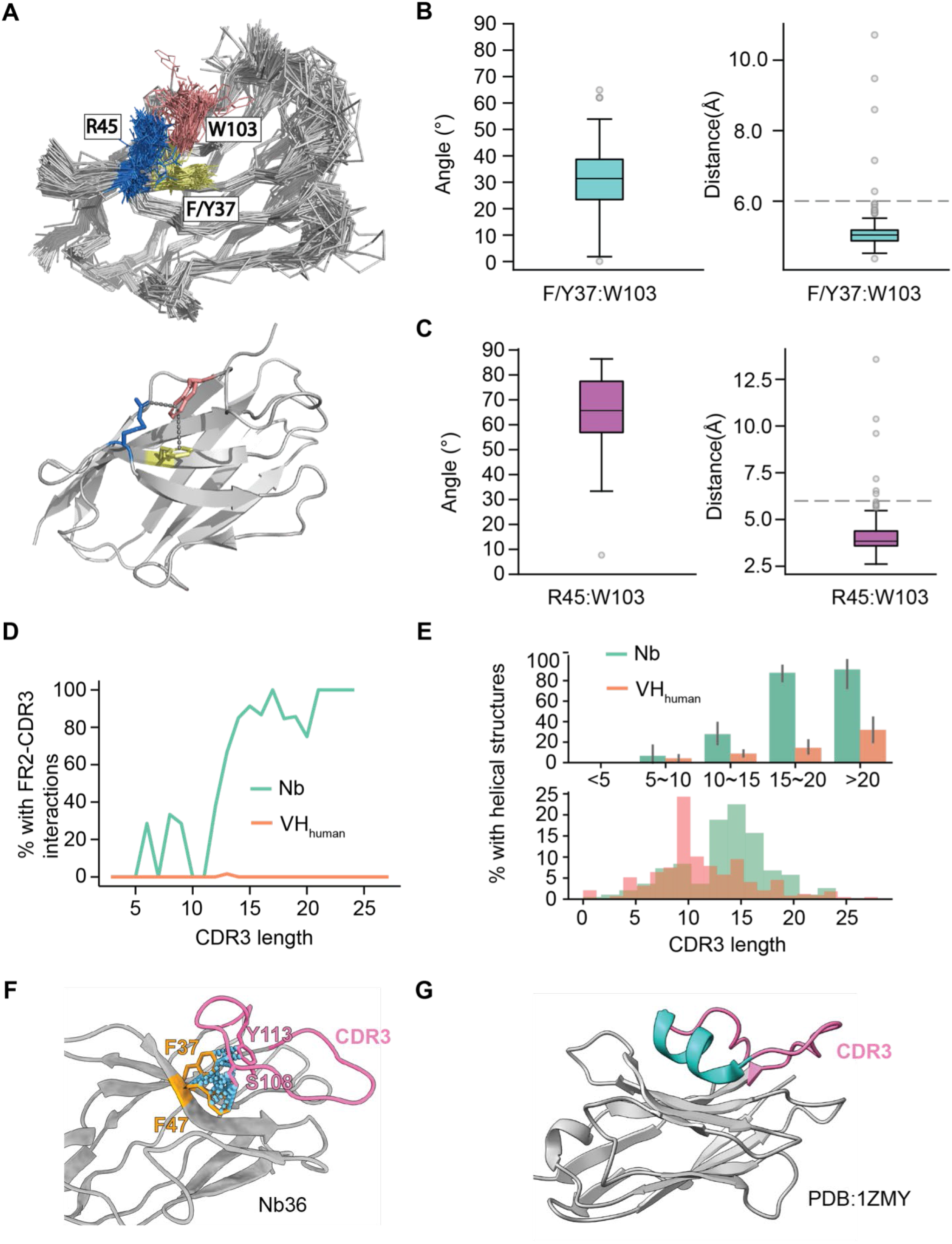
Conserved intramolecular interactions specific to Nbs. **A**. Upper panel: superimposition of 190 Nb structure backbone tracings. F/Y37 (FR2) isin yellow, R45 (FR2) in blue, and W103 (FR4) in salmon. Sidechains of the above residues are shown. Lower panel: the conserved interactions between FR2 and FR4 of a representative Nb (PDB:7JVB). **B**. The distributions of distances and angles of the R45:W103 interactions based on the available Nb structures. **C**. The distributions of distances and angles of the F/Y37:W103 interactions. **D**. Percentage of contacts between 4 FR2 hallmark residues and CDR3 as a function of CDR3 length for Nbs and VH_human_. **E**. Percentage of CDR3s containing additional helix structure in different CDR3 length groups as contrasted to the distribution of CDR3 length for Nbs and VH_human_. **F**. A representative Nb structure showing interactions between a long CDR3 and two conserved FR2 residues (F37 and F47). **G**. A representative Nb structure showing the presence of a short helicase structure in the CDR3 loop.

**Figure 5.**
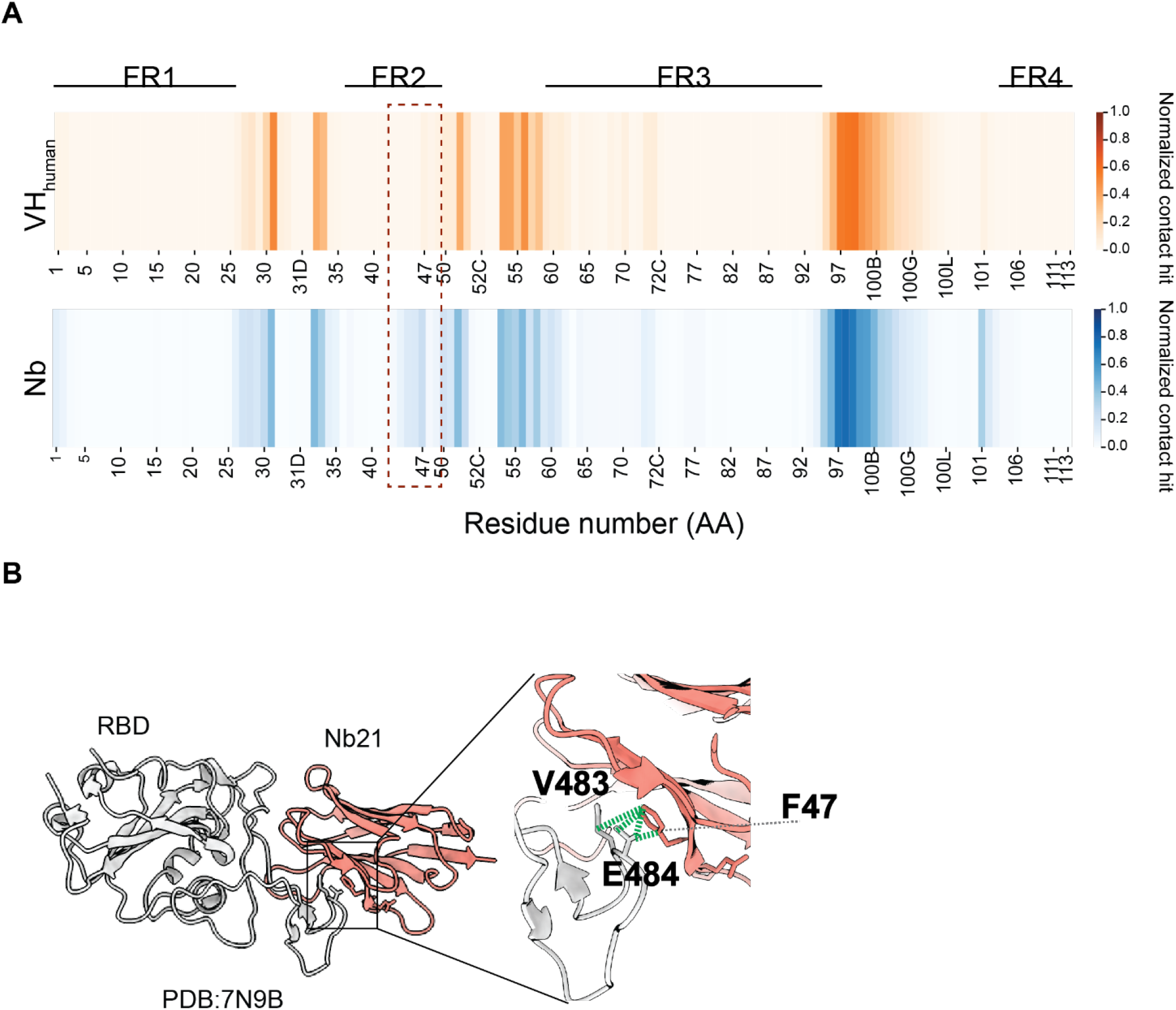
Involvement of Nb FR2 to antigen binding. **A**. Heatmaps showing antigen contact propensity of human VHs and Nbs. Boxed area showed FR2 of Nbs involved more in antigen engagement. **B**. A representative antigen:Nb structure shows the interactions between V483,E484(RBD) and F47(Nb FR2).

Moreover, specific intramolecular disulfide bonds (between CDR3 and CDR1/2) were identified in approximately 10% of Nbs that we have analyzed but were absent in VH_human_ (**Figure S3B**). The disulfide bridges (**Figure S2B**) are exclusively buried for folding and the corresponding residues are not considered for humanization in our software. Finally, intermolecular contact analysis reveals direct involvement of specific FR2 residues (44,45 and 47) for antigen binding. Based on the antigen: Nb structures, approximately 20% of Nbs use these FR2 residues for antigen binding. In addition, it appears *in vivo* affinity matured Nbs use them less extensively for binding compared to *in vitro* selected Nbs (5). Without detailed structure information of antigen: Nb interactions, these FR2 residues should be ideally excluded for humanization to better preserve structural integrity.

### Development of an open-source to facilitate rational Nb humanization

Based on big data analysis, we developed an open-source software to facilitate automated, structure-guided, and robust Nb humanization. Llamanade is composed of five main modules (**Figure 6**): 1) Nb structural modeling, 2) sequence annotation, 3) sequence analysis, 4) structural analysis, 5) humanization score. The input is a Nb sequence in a simple fasta format. The structural model is generated by Modeller (28). Alternatively, the input can be high-resolution structures of Nbs or antigen: Nb complexes. Nb sequence will be annotated using the Martin scheme to define FRs and CDRs. To select the best VH_human_ templates for Nb humanization, we first use Nb sequences (from the NGS database) to fetch the top-20 VH_human_ in our human IgG library that are highly similar to Nbs based on T20_F_ score (**Figure 2**, Methods). 6,070 VHs were obtained after this analysis. The sequences are aligned to calculate the amino acid frequency at each position to generate a position-dependent residue frequency matrix. Annotated input Nbs will be compared to this matrix to select framework residues for humanization. Candidate residues will be selected for humanization if their occurrences are sufficiently low (< 10%) at the corresponding position of VH_human_. Substitution of these residues may result in incompatibility of these rare VH_human_ frameworks with specific CDR loop conformations that compromise the structural and physicochemical properties of the Nbs after humanization.

**Figure 6.**
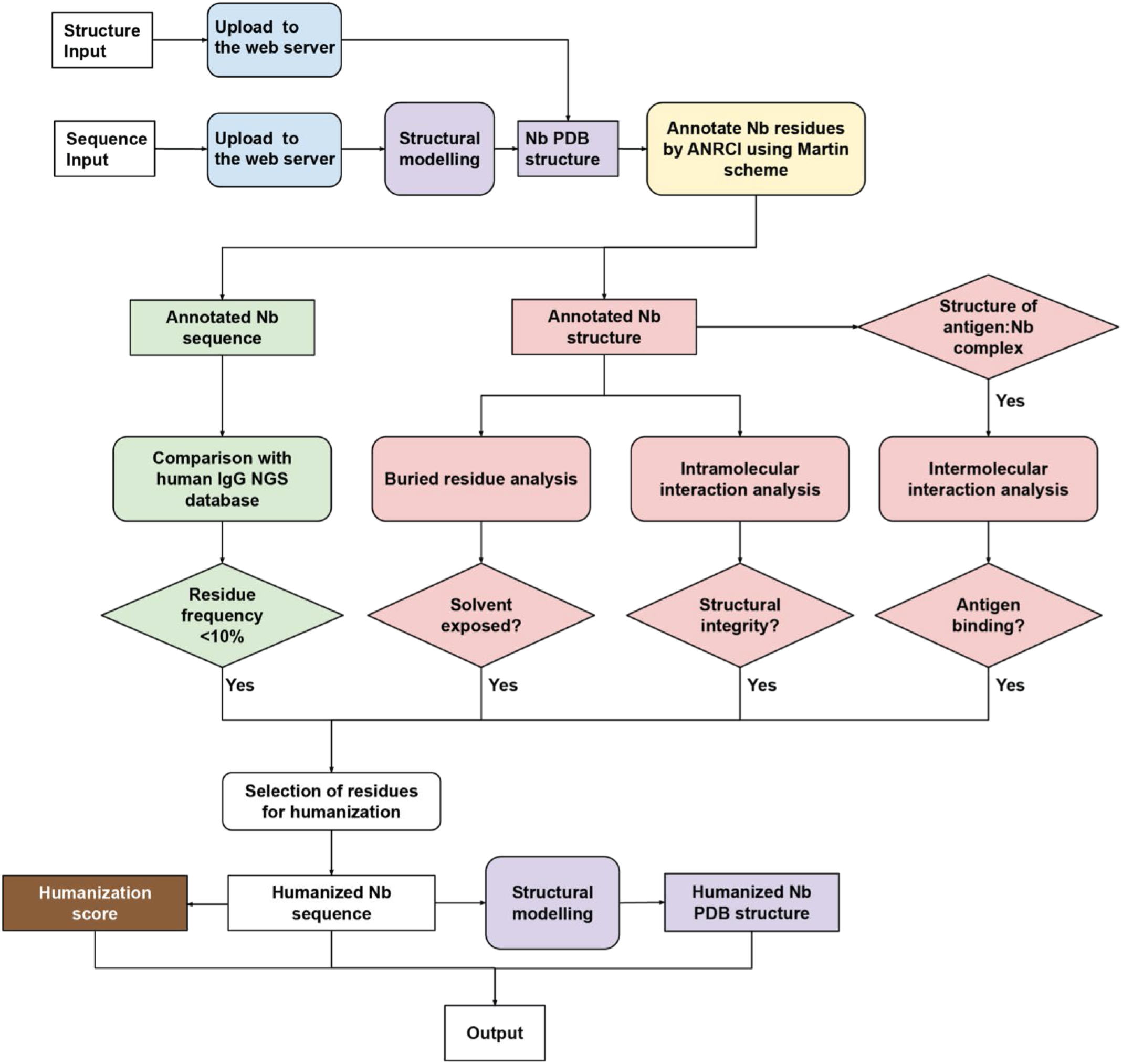
Schematic pipeline of Llamanade. The software is composed of five main modules: 1) structural modeling (purple), 2) sequence annotation (yellow), 3) sequence analysis (green), 4) structural analysis (pink), and 5) humanization score (brown). The input is Nb sequence information in a fasta format. Provided availability, additional structural input can be uploaded to facilitate the analysis. Nb sequences and structures will be first annotated by *Llamanade* to define CDRs and FRs. The sequence analysis module then selects candidate residues for humanization by comparing the sequence to the VH_human_ residue frequency matrix. The structural analysis including degree of burial, intra- and intermolecular interactions will be performed to evaluate the feasibility of humanization for candidate residues. The outputs are the humanized Nb sequence, structure and humanization score.

In parallel, our software will perform structural analysis to determine if the candidate Nb residues are solvent-exposed or not. Only solvent-exposed residues that do not participate in backbone intramolecular interactions (structural integrity), and do not impact antigen: Nb interactions (**Figure 4**) will be humanized. In particular, specific FR2 residues at positions 37, 45, and 47 are not recommended to change because of their involvement in either backbone interactions or antigen-binding (**Figure 3, 4**). A humanization score will be calculated based on T20_F_ to quantify the humanness level of Nbs (**Figure 1**). The output files include the annotated humanized Nb sequence, modeled structure, and the humanization score. *Llamanade* is user-friendly and the full analysis takes less than 1 minute/Nb to complete on a local device. In addition, the service will be deployed to a webserver to provide better accessibility and visualization to the community.

### Humanization of ultrapotent SARS-CoV-2 neutralizing Nbs by Llamanade

To demonstrate the robustness of *Llamanade*, we humanized nine highly potent SARS-CoV-2 neutralizing Nbs that have been recently identified (8). These Nbs target the receptor binding domain (RBD) of the virus spike and are structurally diverse falling into three main epitope classes (5) (**Figure 7A**). Class I Nbs include Nb 21 and presumably Nb64. They can directly block the ACE2 receptor binding sites of RBD to potently neutralize SARS-CoV-2 at as low as below 0.1 ng/ml, which is unprecedented for antiviral antibody fragments (8). Class II Nbs (34, 95, 105, and presumably 93) strongly bind conserved RBD epitopes and neutralize the virus at 30 ng/ml. Class III Nbs (17, 36, and presumably 9) recognize relatively conserved sites including cryptic neutralizing epitopes where large human antibodies may not be able to access. Here robust humanization will help preclinical and clinical development of antiviral Nbs.

**Figure 7.**
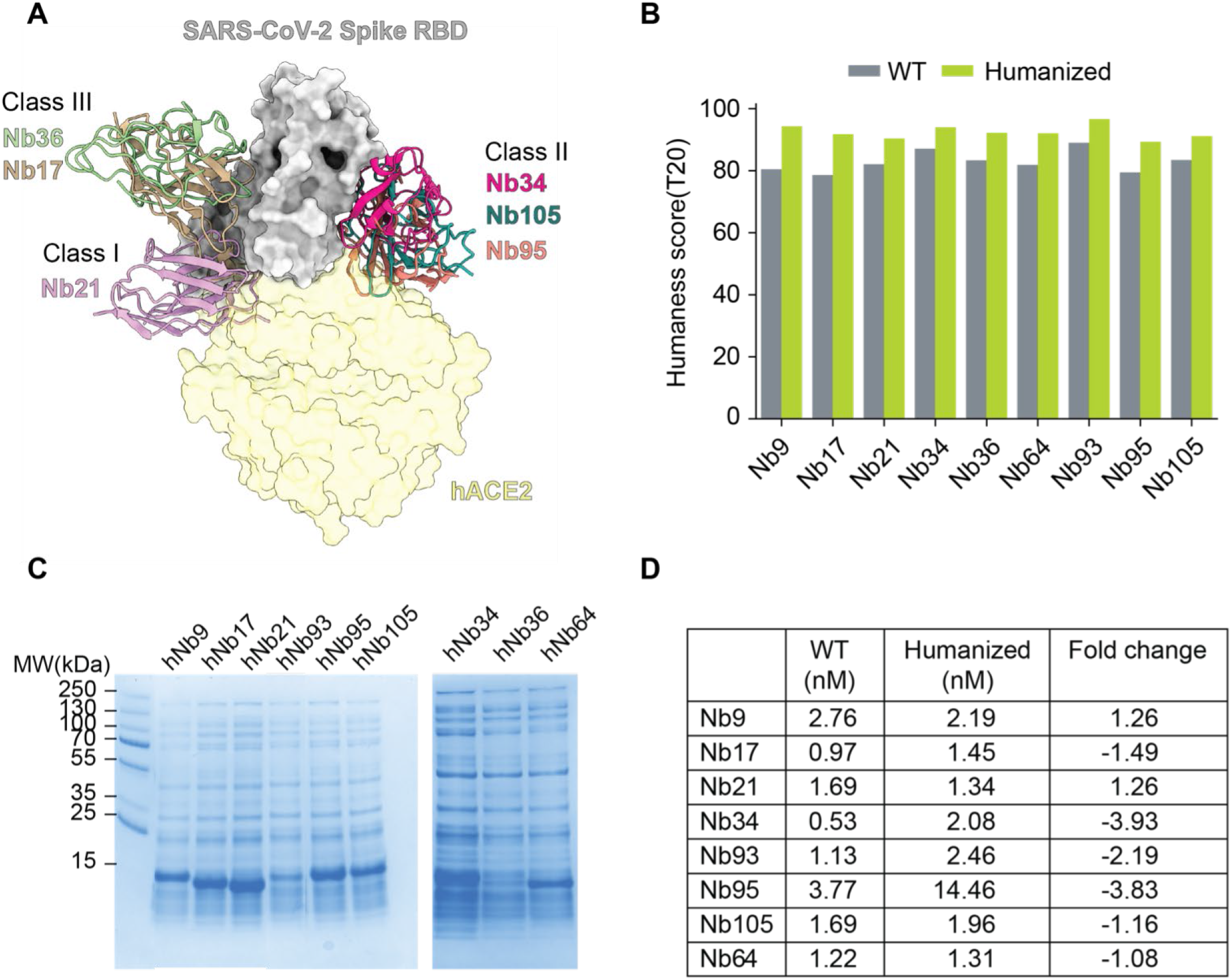
The humanization of SARS-Cov-2 Nbs by Llamanade and verification. **A**. Structure model of binding of three classes of neutralizing Nbs to the RBD of SARS-CoV-2. The angiotensin-converting enzyme 2 (hACE2 in light yellow) is superimposed into the model. **B**. Bar plot comparison of humanness scores (T20) of Nbs before and after humanization. **C**. SDS PAGE gel picture showing the expression level of humanized Nbs. **D**. The relative RBD binding affinities (nM) of wild type (WT) and humanized Nbs measured by ELISA.

Using Nb sequence data as input, these RBD Nbs were humanized by *Llamanade. Llamanade* significantly improved the humanness level of Nb frameworks. The median T20 score of Nbs increased from 82.8 (before humanization) to 92.4 (after humanization), which corresponds to the humanization of seven Nb FR residues (**Figure 7B, Figure S3**). The median framework identity of these highly humanized Nbs to the top matched VHhuman sequence ranges between 90%-95% (**Figure S4**).

After humanization, Nbs were back translated into DNA sequences, which were synthesized *in vitro* and cloned into an expression vector (pET-21b) for recombinant protein productions in *E*.*coli* (**Methods**). 8/9 humanized Nbs were readily expressed in the whole-cell lysis (**Figure 7C**) comparable to their native forms (8). They were one-step purified by His-cobalt resin with excellent solubility and yield (**Methods, Figure S5**). The only exception was Nb36. While hNb36 can still be purified from the soluble cell lysis (**Figure S5**), the majority of the humanized Nb was found in the inclusion body and was excluded for analysis due to potentially inferior solubility. To assess the bioactivities of these humanized Nbs, we performed the enzyme-linked immunosorbent assay (ELISA) and confirmed the comparably high activities (± 4 fold) to the non-humanized precursors (**Figure 7D, Figure S6**). These highly humanized ultrapotent SARS-CoV-2 Nbs recapitulated robust physicochemical and structural properties that are critical for inexpensive manufacturing towards clinical development.

## Discussion

Nbs are characterized by small size, high solubility and stability for advanced biomedical uses. Technology advancement has recently enabled rapid discovery and characterization of tens of thousands of high-affinity and multi-epitope Nbs for specific antigen binding, which opens exciting possibilities for drug development (19). Recently, ultrapotent Nbs have shown great promise as cost-effective antiviral agents to help curve the pandemic caused by SARS-CoV-2. Stable and ultrapotent Nbs can be inhale delivered by aerosolization with high bioavailability to treat pulmonary infections efficiently (11).

Humanization has been considered as a key step for therapeutic development of xeno-species antibodies including Nbs of camelid origin. However, systematic investigations into Nb humanization based on large-scale sequence and structural analyses remain unavailable. To our knowledge, there is no dedicated software available for Nb humanization. To fill this gap, we have developed an open-source *Llamanade* to facilitate robust and fast Nb humanization. Using *Llamanade*, we quickly humanized a cohort of structurally diverse and ultrapotent SARS-CoV-2 neutralizing Nbs. Critically, these highly humanized Nbs demonstrated robust physicochemical and structural properties and high bioactivities comparable to the natural, affinity matured presusors.

The underlying mechanisms of ADA remain to be investigated. However, we note that multiple factors, beyond the use of non-human antibodies, may contribute to the ADA responses. These include: 1) impurity of antibodies, 2) the route and dose of administration, and 3) solubility of antibody or antigen-antibody complex. Inspired by the FDA approval of the first Nb therapy(29), a number of preclinical and clinical programs have been initiated providing critical insights into the safety profiles, dosing regime and efficacy of therapeutic Nbs. For example, recent studies based on dozens of preclinical results found that ADA responses were minor, and were generally non-neutralizing to compromise efficacy (30). Clinical trials have further revealed limited safety issues of Nbs (mostly humanized Nbs) in both patients and healthy volunteers. Nevertheless, ADA has been detected in both preclinical and clinical development. For example, a humanized bivalent anti-cancer Nb targeting human cytokine receptor (hIL-6R) was associated with adverse effects during clinical trials and was terminated prematurely (31). Potential immunogenicity and adverse effects of a non-humanized HER2 Nb were also detected in phase I trial, which showed moderate increase over the course of the trial (32). These studies underscore the requirement of comprehensive evaluations of ADA and toxicology studies before moving Nbs into clinical trials. Here, the development of robust humanization methods will help improve the translational potentials of therapeutic Nbs.

In summary, we have developed a robust, fast, and user-friendly tool to facilitate automated Nb humanization. *Llamanade* is the first dedicated software for Nb humanization. It is freely accessible and can be extended for the analysis of other VH-like scaffolds such as shark single-domain antibodies (V_NAR_) (33). In a parallel effort, Tomer et al. developed AI-enabled software to accurately model Nb structures. These tools will be integrated to further advance Nb-based biomedical research and therapeutic development.

## Data and Methods

### Sequence dataset

VH_human_ and VH_mouse_ sequences were downloaded from EMBL-Ig at:http://web.bioinf.org.uk/abs/abybank/emblig/www/emblig. Data were filtered by removal of duplication and incomplete VH sequences, which leads to a final of 22,450 and 10,696 non-redundant and high quality VH sequences.

### Structural dataset

1694 antigen: IgG complex structures and 246 antigen:Nb complex structures were obtained from AbBank(http://www.abybank.org) (20). Structures were annotated based on the Martin scheme. In addition, 7 SARS-CoV-2 RBD:Nb structures were obtained from a recent study by Dapeng et al(5).

### Humanness T20 score

The T20 scorer was implemented according to the method described in (22). 22,450 non-redundant human VH sequences were used to build the blast reference database. To obtain the T20 score of a given VH/V_H_H sequence, the query sequence will be searched against the reference database by blastp. For an input VH_human_, the top 20 matched sequences will be retrieved and corresponding sequence identities will be averaged to calculate the T20 score. For a VH sequence from any other species, the top 20 matched sequences will be retrieved and used to calculate T20 score. In addition to a full sequence-based T20 scorer, we also implemented a FR based T20 scorer. More specifically, framework sequences of human VHs were extracted to construct the reference database and the framework of query sequence would be used to search. In this study, we implemented several T20 scorers for each individual framework region (FR/FR1/FR2/FR3/FR4).

### Antibody alignment and numbering

Antibodies in the sequence and structural dataset were numbered by ANARCI using the Martin numbering scheme (21). After alignment, aligned positions with more than 99% of gap were removed. The frequencies of amino acids at each aligned position were calculated from MSA to generate the amino acid frequency matrix.

### Structural analysis of buried residues

The degree of burial for a residue is quantified by measuring the depth of the side chain below the protein surface using the ResidueDepth module in BioPython. The depth of the residue side chain is calculated by average distance to surface for all atoms. The protein surface was generated by software MSMS. A cutoff of 3.03 A defining the state of burial or exposure to solvent was used according to (24).

### Contact analysis of antigen-antibody interactions

The Cα distance between two residues was calculated for every antibody-antigen residue pair using ProDy(34). A residue pair with Cα distance less than 8 angstrom(35) was considered for interaction.

### Analysis of nanobody intramolecular interaction

Interactions such the salt bridge, cation-π, π-π were investigated. The following criteria were used to predict the presence of a certain interaction:

Salt bridge: The distance between two opposite charge atoms between two residues is less than 4 angstrom.

Cation-π: The distance between the centroid of the aromatic ring from TRP/PHE/TYR and the charged atom from LYS/ARG is less than 6 angstrom.

π-π: The distance between centroids of two aromatic rings from TRP/PHE/TYR is within 4∼6 angstrom.

### ELISA (Enzyme-Linked Immunosorbent Assay)

Antigens (RBD) were coated onto 96-well ELISA plates, with 150 ng of protein per well in the coating buffer (15 mM sodium carbonate, 35mM Sodium Bicarbonate, pH 9.6) at 4°C overnight. The plates were decanted, washed with a buffer (x×1 PBS, 0.05% Tween 20), and blocked for 2 hours at room temperature (1× PBS, 0.05% Tween 20, 5% milk powder). Nanobodies were serially diluted by 5 fold in blocking buffers. Anti-T7 tag HRP-conjugated secondary antibodies were diluted at 1:5000 and incubated at room temperature for 1 hour. Upon washing, samples were further incubated in the dark for 10 minutes with freshly prepared 3,3’,5,5’-Tetramethylbenzidine (TMB) substrate. Upon quenching the reaction with a STOP solution, the plates were measured at wavelengths of 450 nm with background subtraction at 550 nm. The raw data were processed and fitted into the 4PL curve using the Prism Graphpad 9.0. IC50s were calculated and fold changes of binding affinity were calculated to generate the heatmap.

## Author contributions

Y.S. conceived the study. Z.S. analyzed the data and developed the software. Y.X. and Z.S. performed the validation experiments. Y.S. and Z.S. wrote the manuscript with input from Y.X. and I.B.

## Acknowledgement and conflict of interest

Funding: This work was funded by NIH 1R35GM137905-01 (Y.S.), a grant from MJFF and the Alzheimer Disease foundation (Y.S.). The University of Pittsburgh has submitted a provisional patent application related to this work in which Z.S. and Y.S. are co-inventors. We thank Jeff Kim for proof-reading of the manuscript.

