## Supplemental information for "*Llamanade*: an open-source computational pipeline for robust nanobody humanization"

**Supplementary Information**

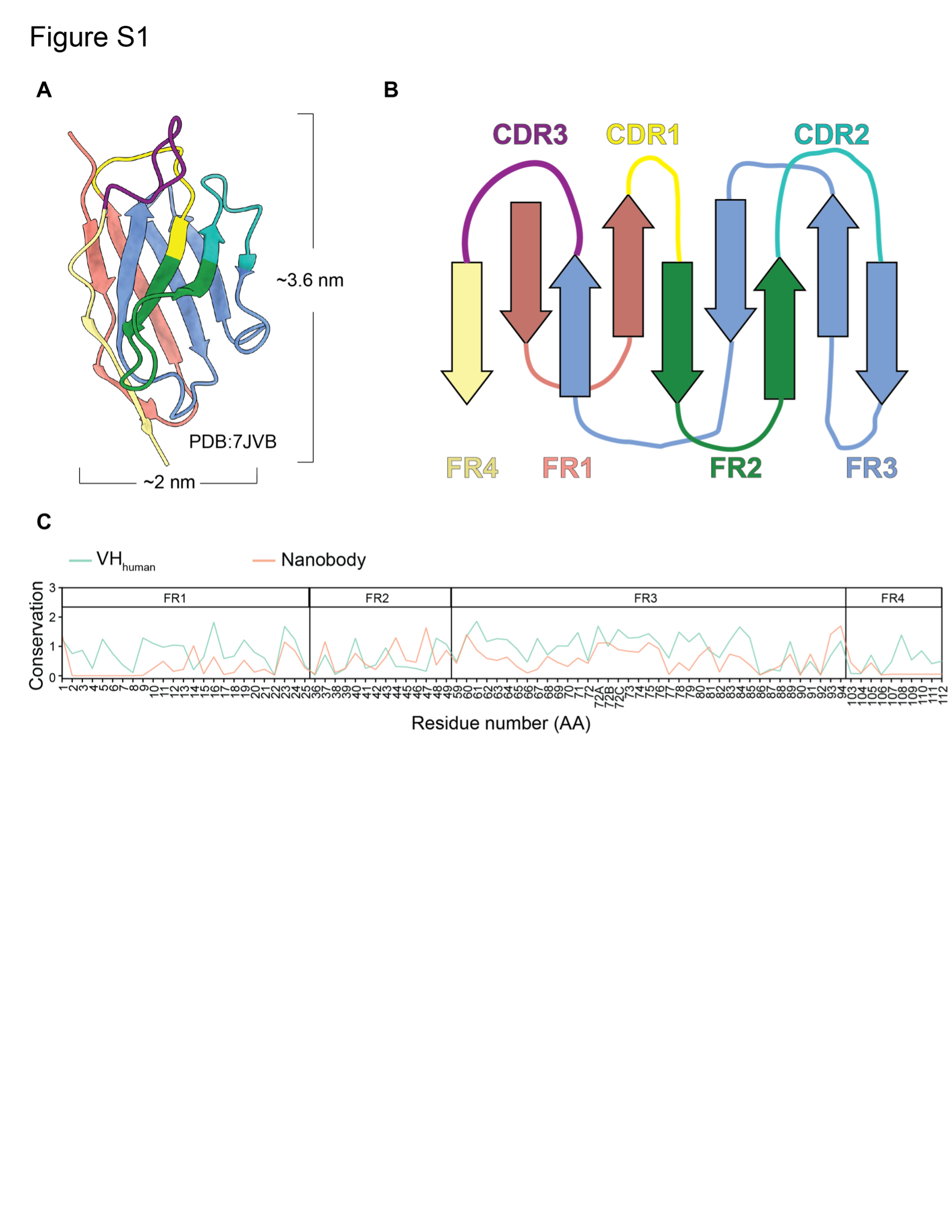

**Figure S1. Quantitative analysis of sequence conservation of Nbs and human VHs**

**A.** Crystallographic structure of a nanobody (PDB: 7JVB). Frameworks (FRs) and complementarity-determining regions (CDRs) are shown in different colors (FR1: salmon; CDR1: yellow; FR2: green; CDR2: cyan; FR3:blue; CDR3: purple; FR4: khaki)

**B.** Schematic of nanobody structure.

**C.** The conservation of the framework residue is calculated based on the entropy of amino acid variations in a given position in multiple alignments.

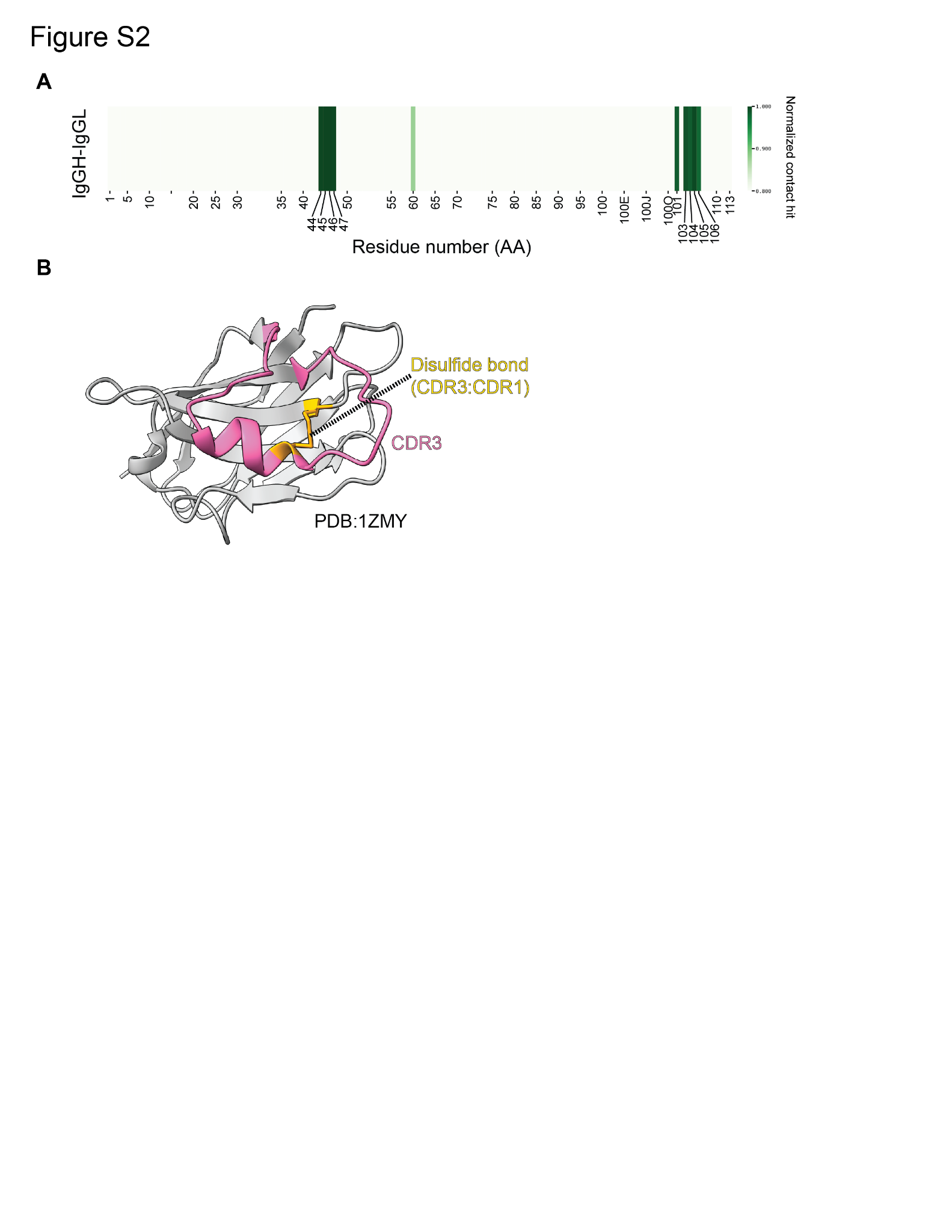

**Figure S2. VH-VL contacts in human IgGs and unique disulfide bond in Nbs**

**A.** Heatmap showing the VH-VL contact propensity of VH_human_.

**B.**  A representative Nb structure (PDB: 1ZMY) showing the disulfide bond formed between CDR3 and CDR1.

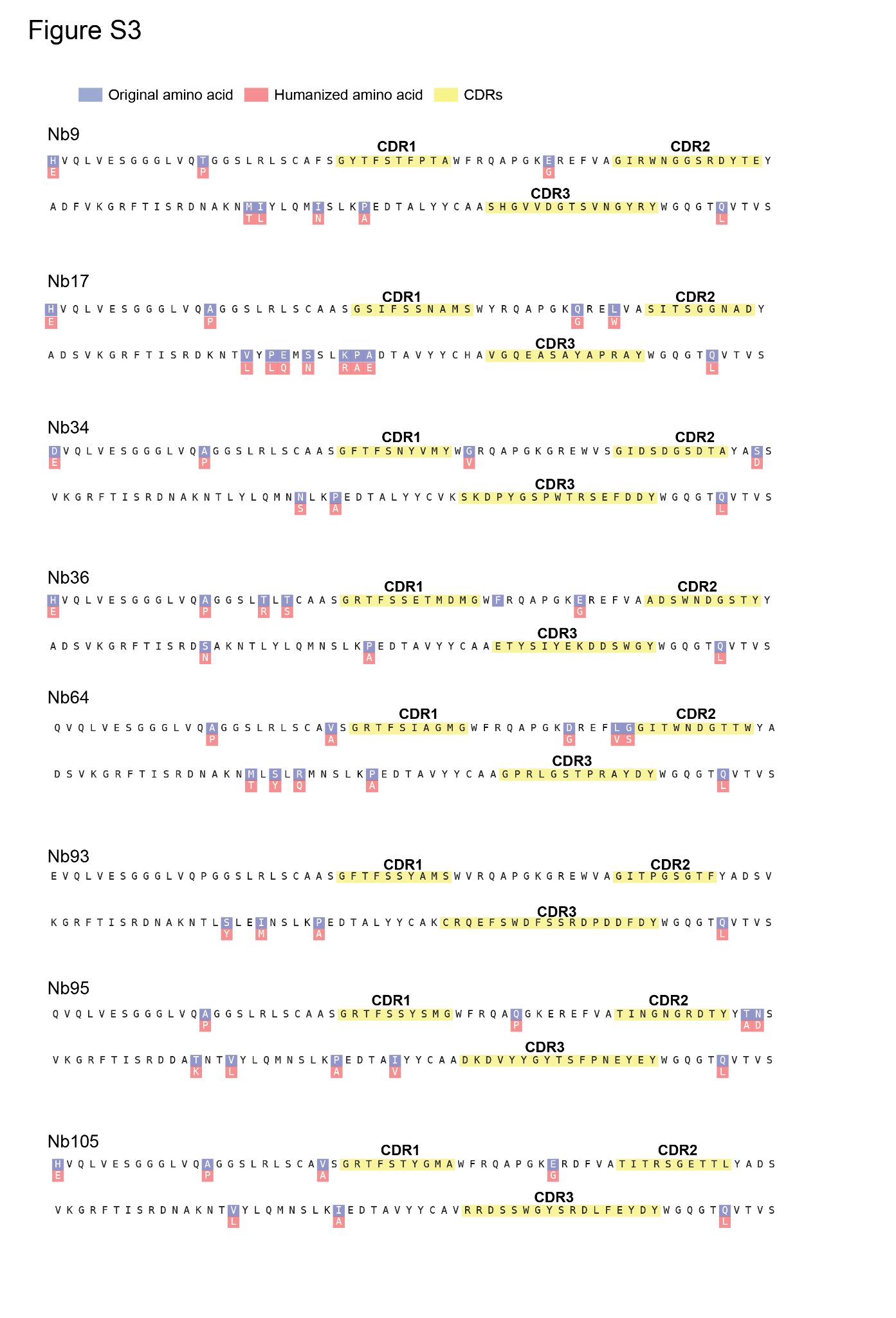

**Figure S3. Sequence alignments of wild type and humanized Nbs.**

CDR sequences were highlighted. Humanized residues were in blue (prior) and red (after humanization).

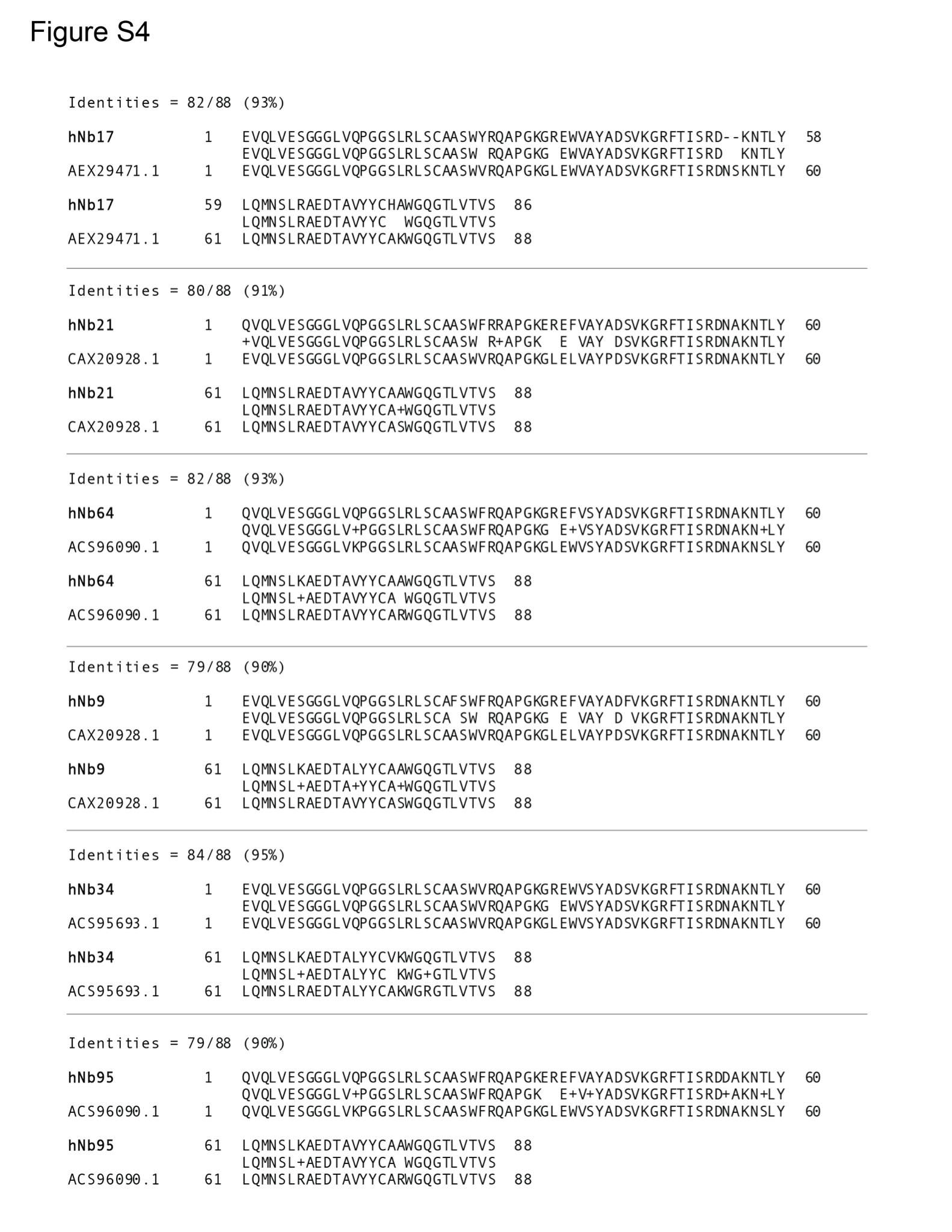

**Figure S4. Framework sequence alignments of humanized Nbs to the best matched VH_human_**

Framework sequences of humanized Nbs are used to search the VH_human_ framework sequences with highest sequence identity.

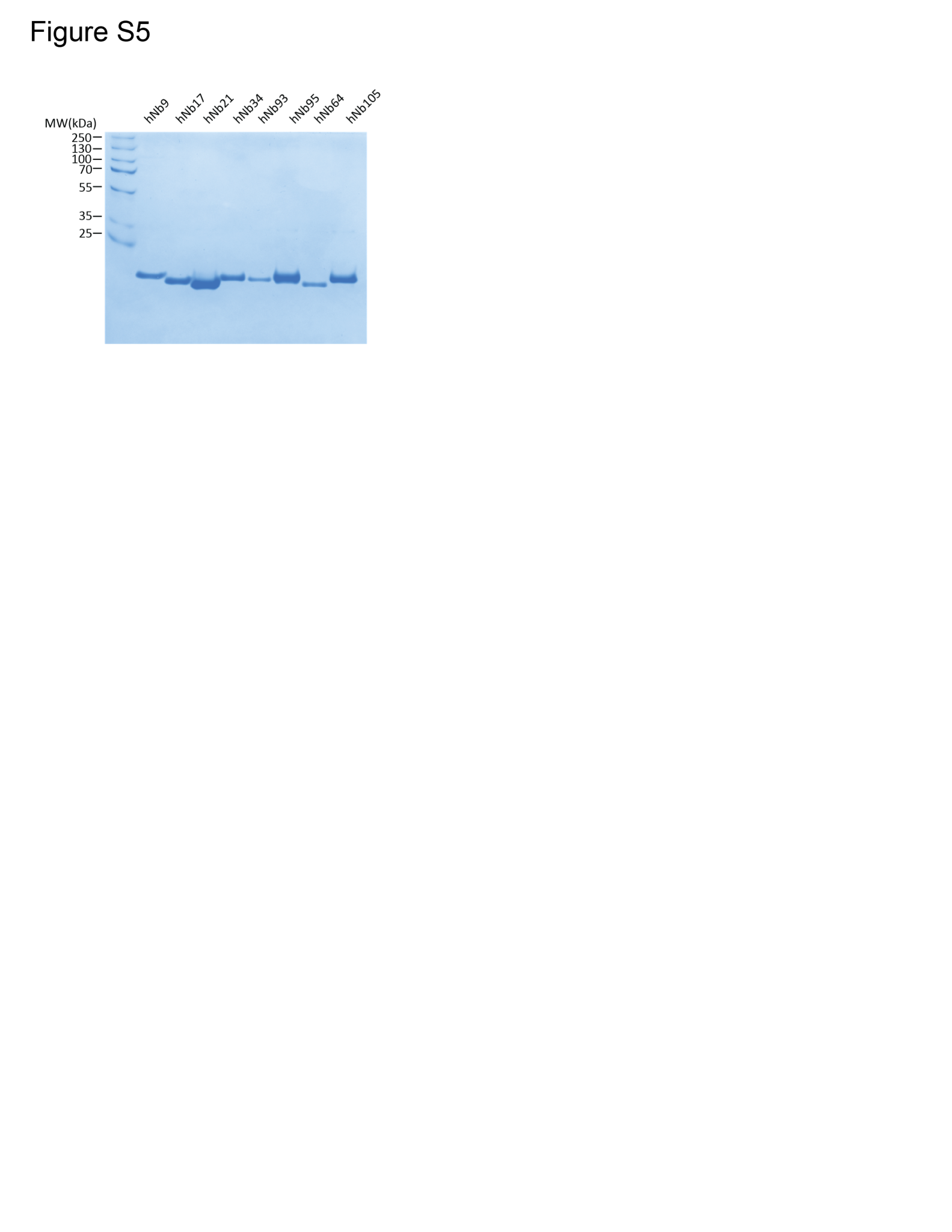

**Figure S5. One-step purification of humanized SARS-CoV-2 Nbs.**

Humanized Nbs were purified from E.coli whole cell lysis by using His6-cobalt resin. After imidazole elution, highly purified Nbs were analyzed by SDS-PAGE.

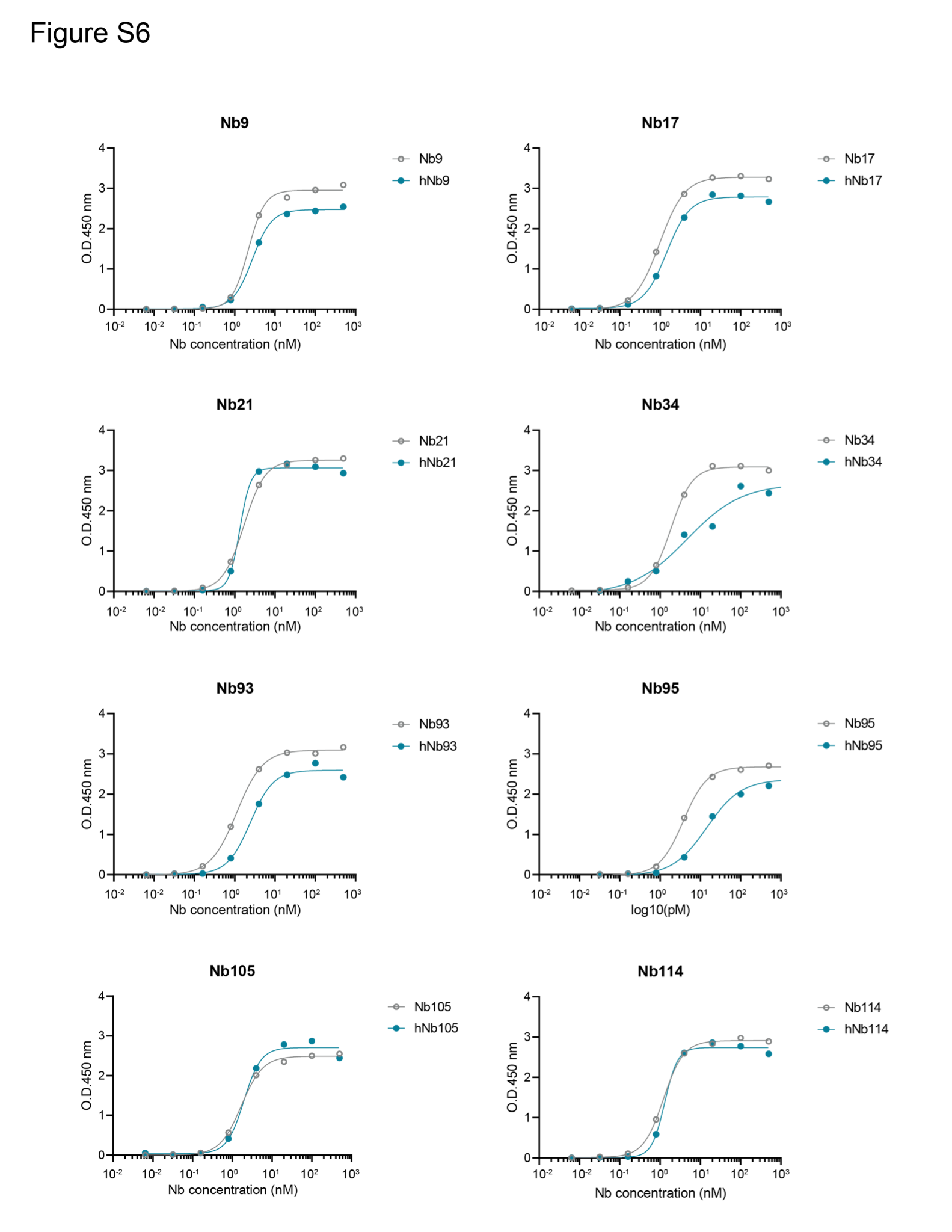

**Figure S6. RBD (SARS-CoV-2 spike) binding of wild-type and humanized Nbs by ELISA.**

The ELISA Optical Density readings at 450 nm were plotted against Nb concentrations (nM). hNb: humanized Nb.
